# Quantifying the Environmental Impact of Cell Culture

**DOI:** 10.64898/2026.05.21.720586

**Authors:** Isobel Taylor-Hearn, Jack Llewellyn, Charlotte E. L. Mellor, Martin Farley

**Author notes:** Correspondence should be addressed to M.F.

## Abstract

Laboratory research generates substantial plastic waste and associated greenhouse gas emissions, yet researchers often lack practical tools for quantifying the environmental impact of routine protocols or identifying realistic opportunities for reduction. Here, we present an open-source calculator for estimating plastic use and carbon dioxide equivalent emissions from laboratory protocols, using item weight, plastic composition, and estimated cradle-to-grave carbon footprint factors. We apply the tool to a standard cell culture workflow to demonstrate how evidence-based protocol adjustments can reduce plastic consumption and emissions without affecting experimental design or efficiency. The calculator is designed to be transparent, adaptable, and extendable, allowing researchers to add new consumables and tailor analyses to their own laboratory practices. This work provides a quantitative framework for translating sustainability principles into measurable, protocol-level changes and supports more environmentally responsible decision-making in biomedical research.

## INTRODUCTION

Laboratory research is resource-intensive and contributes disproportionately to global waste generation. Scientific laboratories have been estimated to produce around 5.5 million tonnes of plastic waste annually—approximately 2% of all plastic waste—despite researchers representing only 0.1% of the global population (Schneegans et al., 2021; Urbina et al., 2015). Although the methodology underlying the 2015 estimate is unclear, more recent data from a survey of seven research laboratories suggest plastic waste generation of 30–240 kg per researcher per year (Weber et al., 2025). Even the lower end of this range exceeds average per-capita plastic consumption globally, and it excludes plastic use outside laboratory settings. Notably, laboratories performing cell culture produced the highest waste volumes (Weber et al., 2025), reflecting the heavy dependency of these workflows on single-use plastics and sterile consumables.

Cell culture remains a cornerstone of modern life sciences, underpinning critical advances in cancer biology, immunology, and regenerative medicine. However, it is also a resource-intensive technique, often generating significant volumes of plastic waste and consuming substantial energy and reagents. Without conscious management, routine cell culture can have a considerable, and often overlooked, environmental footprint.

To address these challenges, structured frameworks such as the Laboratory Efficiency Assessment Framework (LEAF) have been developed to promote sustainability in wet laboratory environments. LEAF provides standardised actions and recognises laboratories that adopt practices to reduce waste and energy use. Through laboratory sustainability audits conducted at The University of Manchester, we identified numerous simple yet effective interventions that reduced the environmental impact of cell culture without compromising experimental quality or reproducibility. However, comparing such interventions in a robust and transparent way required a consistent method for quantifying their effects.

To address this, we developed an online environmental impact calculator that enables protocol-level quantification of plastic waste and associated carbon footprint under different use and disposal scenarios. The calculator provides a standardised framework for evaluating the consequences of workflow changes and allows direct comparison between conventional and modified laboratory practices.

In this study we present a series of sustainable cell culture practices identified through laboratory audits. We use the online tool to quantify the environmental impact of each intervention to demonstrate that small procedural adjustments, when applied to high-frequency routines, can generate substantial cumulative reductions in plastic waste and greenhouse gas emissions. We provide an updated “green” cell culture protocol that integrates the proposed interventions into an adaptable framework, directly comparable to a conventional baseline workflow. This protocol is designed to be flexible, enabling researchers to tailor sustainability improvements to their specific cell types, culture formats, and experimental requirements.

The calculator is made available as a complementary web-based resource and is designed to be reusable and extendable across laboratory contexts. Integrated within SPARKHub, an open-access platform developed to support greener research practices, it provides an accessible implementation of the quantification approach used throughout this study. Together, the tool and the cell culture case study provide both a practical and transparent framework for environmental assessment and a set of evidence-based examples showing how routine laboratory workflows can be redesigned to reduce their environmental footprint without compromising experimental integrity.

## METHODS

An online calculator was developed to enable protocol-level environmental impact quantification (*Environmental Impact Calculator*, 2026). A consumable dataset was generated by weighing a range of commonly used laboratory plastic items and assigning each item a primary polymer type based on manufacturer-provided material specification. Measured consumable weights were linked to published cradle-to-grave carbon footprint assessment (CFA) values for major plastic classes and disposal pathways (Ragazzi et al., 2023). This enabled estimation of the environmental impact of plasticware over its full lifecycle, including raw material extraction, manufacturing, transport, distribution, use, and various end-of-life disposal routes.

Where equivalent products were available from multiple suppliers, both brand-specific weights and a mean item weight were recorded. Mean values were used in subsequent analyses to provide representative estimates and minimise bias towards any single manufacturer. The analysis therefore focused on workflow modifications, such as reducing consumable use or changing disposal practice, rather than on substitution between suppliers. A full online dataset of consumables, weights, and polymer classifications is provided alongside the calculator.

For clarity and consistency, packaging weight was not included in subsequent quantifications, as various packaging options were available for each consumable. However, packaging data can be incorporated into the calculator, and researchers may achieve additional reductions beyond those quantified here by selecting lower-packaging options such as unracked or multipack formats.

Four disposal pathways were included in the analysis: (i) no disposal (reuse); (ii) incineration with energy recovery; (iii) autoclaving followed by incineration with energy recovery; and (iv) clinical waste disposal via high-temperature incineration. This framework enabled direct comparison between consumable reduction strategies and improved waste routing practices, enabling a consistent basis for estimating environmental savings.

To demonstrate the application of this framework in a biologically relevant setting, we modelled a representative cell culture workflow involving a single researcher maintaining two adherent cell lines, each passaged twice weekly. This corresponded to four passages per week, or 752 passages annually assuming a 48-week working year. This scenario provided a standardised baseline for evaluating how specific workflow modifications alter plastic use and carbon footprint.

The calculator was designed as an extendable resource, allowing users to contribute additional consumables and expand the dataset over time. It is therefore presented as a practical implementation of the quantification framework used in this study and as a reusable resource for protocol-specific environmental assessment in other laboratory workflows.

## RESULTS

In this study, we compare a conventional cell culture workflow with an updated “green” protocol that incorporates the sustainability swaps outlined in this paper. **Figure 1A** illustrates the baseline workflow, representing a researcher passaging two adherent cell lines in a single session. This protocol reflects common practice within our institution and is not intended as a “worst-case” scenario. Rather, it represents a reasonable and widely used approach, assuming competent sterile technique (for example re-using a serological pipette across multiple flasks without contacting the necks to avoid contamination).

**Figure 1.**
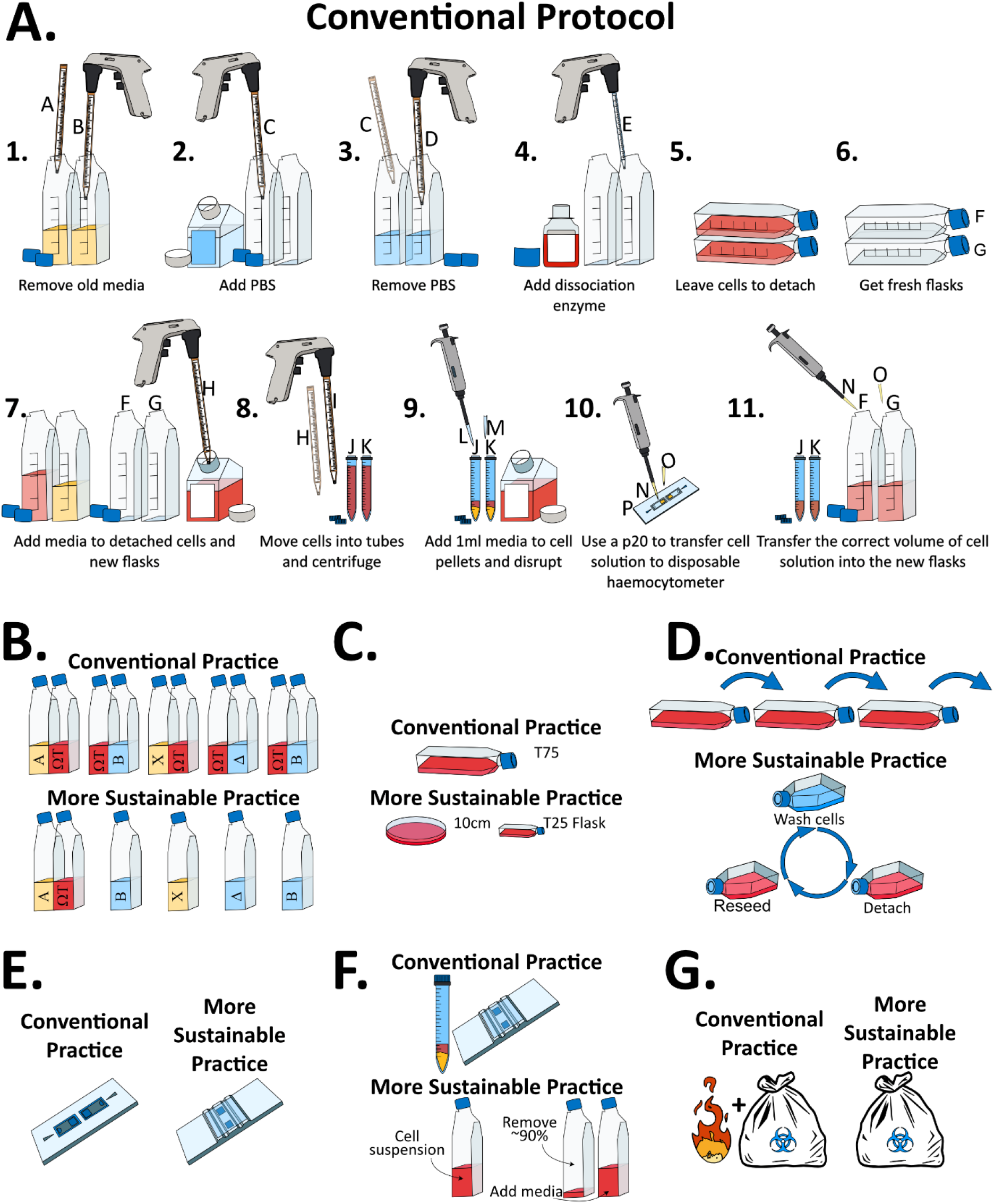
**A:** Conventional cell culture workflow, assuming a researcher passages two cell lines in one session with good sterile technique. This baseline approach generates approximately 169 g of plastic waste per session. **B:** Swap 1: coordinate cell culture to avoid duplication. If all researchers are mainingaing a reference cell line, e.g. ΩT in the illustration), they can coordinate culture lines so that only a single culture is maintained at any given time. **C:** Swap 2: select culture vessels appropriate to experimental needs. T75 flasks can be replaced with 10 cm dishes to reduce waste but maintain culture area. T25 flasks can be used for routine passaging, where only small cell numbers are required. **D:** Swap 3: passage cells directly within their existing container. After detachment, the cell suspension is diluted with fresh medium and a portion of the cell suspension is kept in the original culture vessel to reseed. **E:** Swap 4: replace single-use haemocytometers with reusable alternatives, e.g. reusable glass counters or automated counters with a reusable slide. **F:** Swap 5: passage cells ratiometrically to minimize consumable use. Passaging cells ratiometrically instead of centrifuging and counting avoids the use of a 15 ml tube, filter tip, and any single-use haemocytometers. **G:** Swap 6: dispose of cell culture plastics through the most suitable waste stream. Deactivating biological waste with chemical decontamination allows waste to be sent directly for incineration with energy recovery, instead of requiring an autoclaving step.

**Figure 1.**
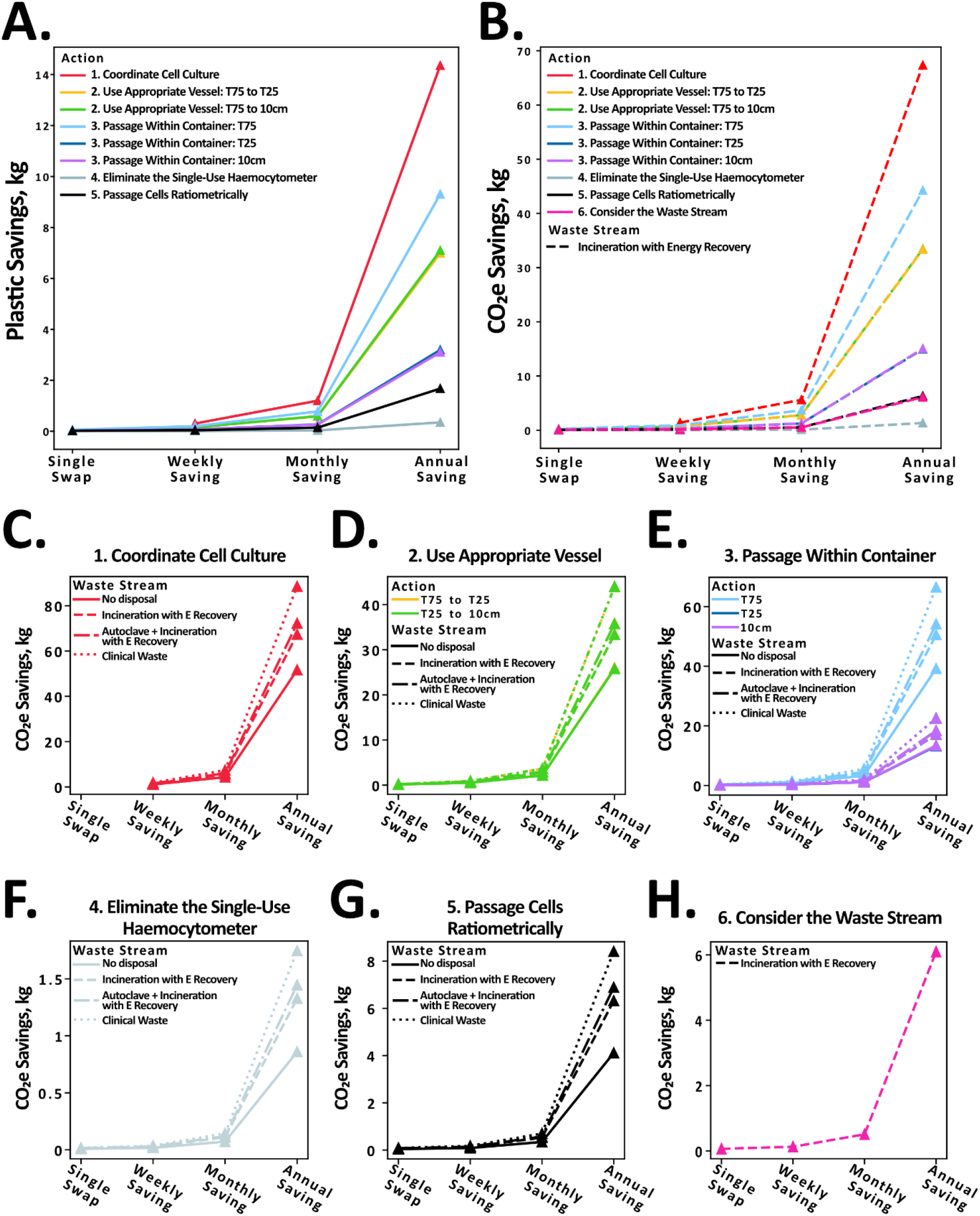
**A-B:** Plastic (**A**) and CO_2_e (**B**) savings associated with the swaps set out in this paper, assuming disposal via incineration with energy recovery before and after the swap. Further swap details and full quantification can be found in **Table 2**. Full quantification is set out in **Table 2** (plastic savings) and **Table 3** (CO_2_e savings). All swaps assume a researcher passaging two cell lines, each twice per week, over a 48-week working year. **C-H:** CO_2_e savings from Figure 2B, separated by swap and waste stream. Full quantification is available in **Table 3**. Swaps are plotted simultaneously to allow readers to see the relative savings associated with each swap. Swaps 1-5 are all shown for 4 waste streams: (i) No disposal (reuse), (ii) Incineration with energy recovery, (iii) Autoclave followed by incineration with energy recovery, and (iv) Clinical waste disposal via high-temperature incineration. Swap 6 considers deactivation of biological waste with chemical decontamination. This saving therefore considers the energy difference between waste streams (iii) and (ii). Importantly, the large CO_2_e savings observed for swaps involving clinical waste reflect the high energy intensity of this disposal route, not a change in waste stream. For these swaps, clinical waste disposal is assumed both before and after the intervention; the savings arise solely from reducing the amount of material entering the clinical waste stream, not from switching to a lower-energy disposal route. Except for Swap 6, the disposal pathway is identical pre- and post-swap. Additional CO_2_e reductions may be achieved when appropriate by combining material-reduction swaps with a transition to a less hazardous waste stream where appropriate.

**Figure 2.**
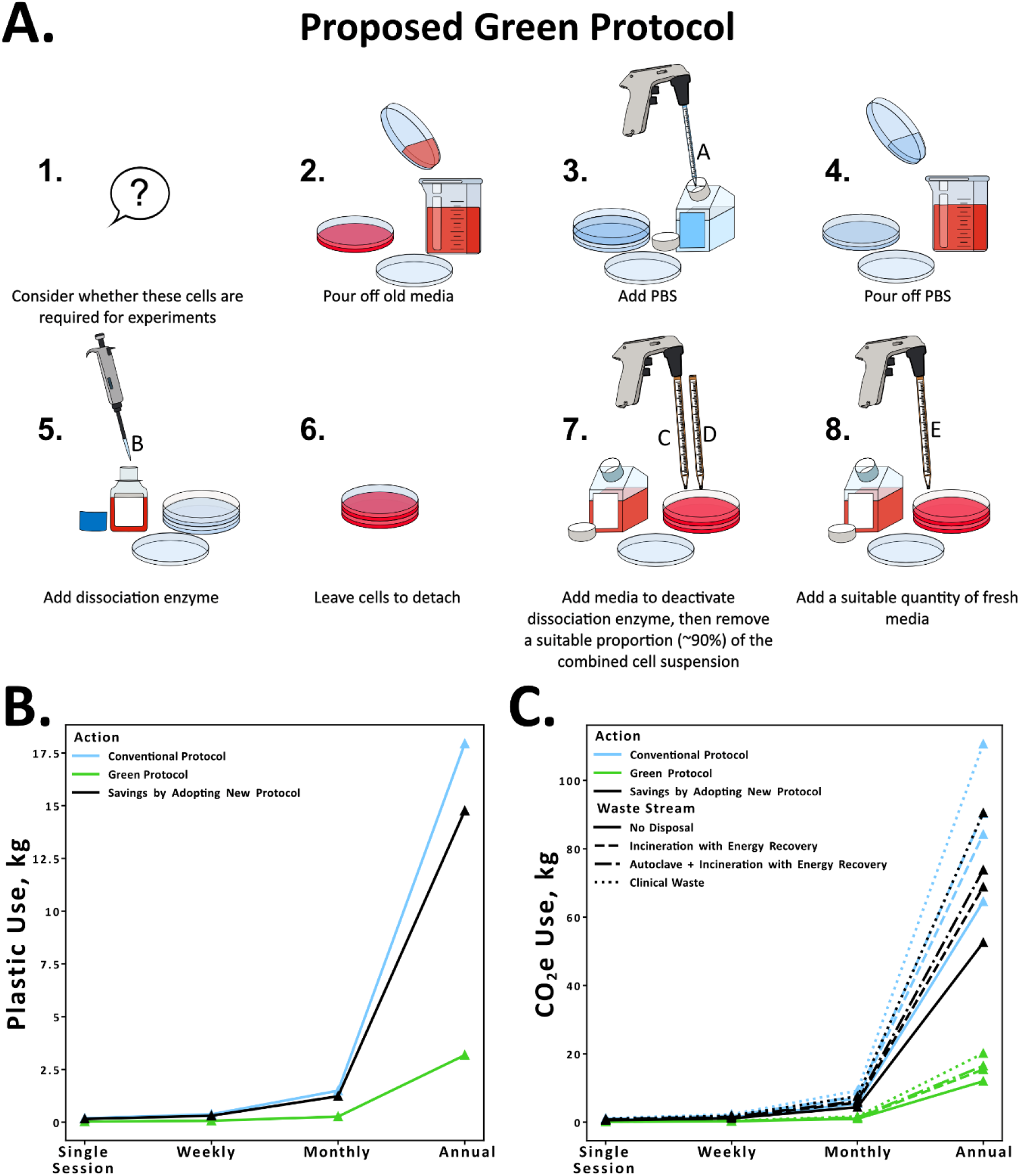
**A:** A “green” cell culture workflow that incorporates the swaps laid out in this paper, assuming a researcher passages two cell lines in one session. This updated protocol generates around 33 g plastic per session (broken down in **Table 5**), a reduction of 154 g per session compared to the standard protocol (**Figure 1A**). **B-C:** Plastic waste (**B**) and CO_2_e (**C**) generated by the conventional protocol (**Figure 1A**) and proposed green protocol (**Figure 3A**). Savings associated with updating the protocol are also illustrated. Full quantification is set out in **Table 4**. Quantification assumes an identical disposal route before and after the protocol update. Accordingly, the large CO_2_e savings observed for clinical waste reflect the high energy intensity of this disposal route, with reductions arising solely from decreasing the quantity of material entering the clinical waste stream, rather than from any change in waste classification. Additional CO_2_e savings may be achieved where appropriate by combining the updated protocol with diversion to a lower-energy waste stream where appropriate.

This design ensures that any observed reductions in plastic use and environmental impact arise from genuine improvements achievable in routine culture, rather than from artificially inflated baseline waste. The conventional workflow generates approximately 187 g of plastic waste and an embodied carbon footprint of 672 g CO_2_e per session (equivalent to approximately 18 kg plastic and 65kg CO_2_e annually, **Table 1**). Depending on the disposal route, this increases to 877 g CO_2_e (incineration with energy recovery), 940 g CO_2_e (autoclaving prior to incineration with energy recovery), or 1152 g CO_2_e (clinical waste disposal) per session.

**Table 1.** Estimated plastic generated per cell passaging session following the standard protocol (outlined in **Figure 1A**). Calculations assume one researcher passaging two cell lines in a single session. The procedure requires 16 disposable plastic items with a combined mass of 186.8 g, broken down by consumable type as shown. Large weights are shown in red, while smaller weights are shown in green.

| Consumable | Labels on Diagram | Consumable Weight, g | Number Required | Total Weight, g |
| --- | --- | --- | --- | --- |
| 10ml Stereological Pipette | A,B,C,D,H,I | 8.3 | 6 | 49.7 |
| 5ml Stereological Pipette | E | 7.5 | 1 | 7.5 |
| T75 Flask | F,G | 55.4 | 2 | 110.7 |
| 15ml Tube | J,K | 6.6 | 2 | 13.3 |
| P1000 Filter Tip | L,M | 0.8 | 2 | 1.6 |
| P20 Filter Tip | N,O | 0.3 | 2 | 0.6 |
| Disposable Haemocytometer | P | 3.6 | 1 | 3.6 |
| Totals |  |  | 16 | 186.8 |

The remainder of this article outlines practical modifications to the baseline protocol, quantified in terms of both plastic use and cradle-to-grave CO_2_e emissions. These measures are not intended as an all-or-nothing package, but rather as a set of modular interventions that can be adopted individually according to experimental needs. The updated workflow integrates all of the sustainability swaps, demonstrating that routine culture can be made substantially more resource-efficient without compromising sterility, reproducibility, or flexibility across cell types. Importantly, even partial adoption of these measures can yield meaningful reductions in plastic consumption and carbon footprint. We recommend that such practices be evaluated within the context of individual laboratories, discussed collectively to safeguard scientific integrity, and implemented in consultation with local health and safety officers. In-text CO_2_e values are reported assuming waste is autoclaved and subsequently incinerated with energy recovery, reflecting the most common disposal pathway at our institution. However, values for all four waste streams can be found in the accompanying tables.

### Swap 1: Coordinate Cell Culture to Reduce Duplication

A simple but effective step toward sustainability is to culture cells only when necessary. In many cases, it is more efficient to maintain frozen stocks and revive cells as required, rather than continuously culturing multiple flasks. This is particularly relevant for widely used “core” or reference cell lines. Instead of each researcher maintaining their own flask, laboratories can coordinate by rotating responsibility so that only a single culture is maintained at any given time. This simple adjustment substantially reduces media, plasticware use, incubator space, and the associated energy and CO_2_e footprint, without compromising access or cell line integrity. For example, in a group of five researchers each passaging one flask of wildtype cells and one additional line per week, coordination would reduce the number of passages from 20 to 12 per week (**Figure 1B**). As shown in **Table 2**, this consolidation reduces the workload by 8 passages per week, corresponding to 1,496 g of plastic (8 × 187 g) saved across the lab, or an average of 299 g per researcher. Over a 48-week working year, this amounts to more than 14 kg plastic waste and almost 53 kg CO_2_e saved per researcher.

**Table 2.** Estimated plastic savings from alternative cell culture practices. Calculations assume one researcher culturing two cell lines, each passaged twice per week over a 48-week working year. Small plastic savings are highlighted grey while large savings are highlighted in green. **Swap 1:** Coordinated maintenance of a shared reference cell line. Assumes five researchers, each normally passaging two cell lines (one being the reference). Under coordination, only one researcher maintains the reference line at a time, reducing duplication. This eliminates 8 passages per week across the group, saving 1,496 g plastic (8 × 187 g) annually, or an average of 299 g per researcher. **Swap 2:** Use of alternative culture vessels. “Single swap” values indicate the saving from replacing one T75 flask (two flasks are required per session in the hypothetical scenario). **Swap 3:** Passaging within the same dish. The culture vessel is replaced monthly (every eight swaps). “Single swap” values correspond to one culture vessel (two required per session). **Swap 4:** Replacement of disposable haemocytometers with a reusable alternative. Assumes each passage uses one chamber of a two-chamber device. **Swap 5:** Ratiometric passaging, which removes the need for a two 15 ml centrifuge tubes, two p20 filter tip, and a disposable haemocytometer per session (passaging two cell lines).

| Action | Plastic Savings, g |  |  |  |
| --- | --- | --- | --- | --- |
|  | Single Swap | Weekly | Monthly | Annual |
| **Swap 1: Coordinate Cell Culture** | NA | 299 | 1196 | 14346 |
| **Swap 2: Use Appropriate Vessel: T75 to T25** | 36 | 146 | 582 | 6985 |
| **Swap 2: Use Appropriate Vessel: T75 to 10cm** | 37 | 148 | 591 | 7092 |
| **Swap 3: Passage Within Container: T75** | 55 | 194 | 775 | 9299 |
| **Swap 3: Passage Within Container: T25** | 19 | 66 | 266 | 3187 |
| **Swap 3: Passage Within Container: 10cm** | 18 | 64 | 258 | 3093 |
| **Swap 4: Eliminate the Single-Use Haemocytometer** | 4 | 7 | 29 | 344 |
| **Swap 5: Passage Cells Ratiometrically** | 17 | 35 | 139 | 1670 |

**Table 3:**
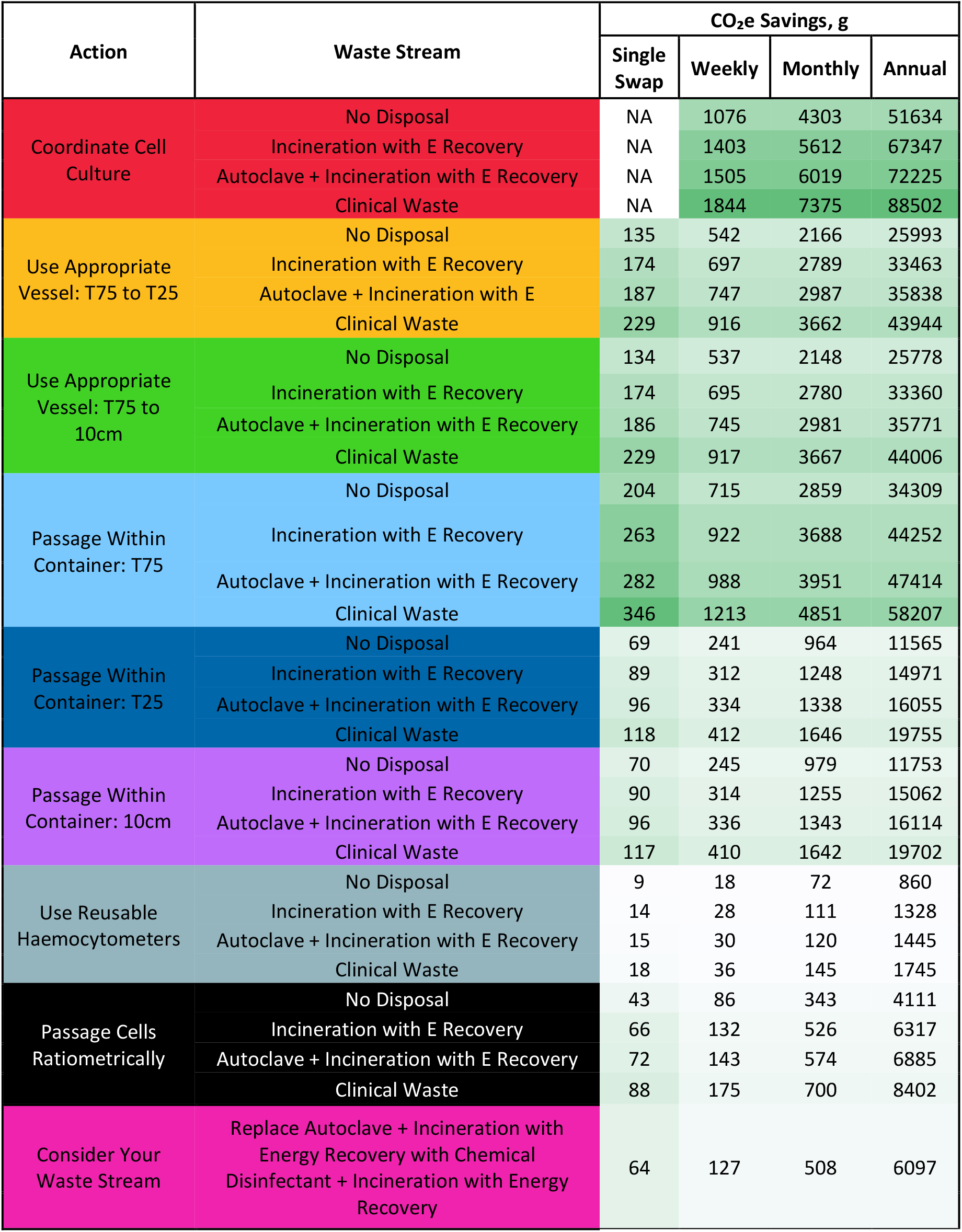
Estimated CO_2_e savings corresponding to the plastic reductions described in **Table 2**, calculated under four different disposal routes: (i) no disposal, (ii) incineration with energy recovery, (iii) autoclaving followed by incineration with energy recovery, and (iv) high-temperature clinical waste incineration. Calculations assume one researcher culturing two cell lines, each passaged twice per week over a 48-week working year. “Consider your waste stream” does not include life cycle assessments of the disinfectant chemicals, nor the additional energy required for ancillary processed (e.g. heating water for washing). Smaller CO_2_e savings are highlighted grey while larger savings are highlighted in green.

**Table 4:** Estimated CO_2_e and plastic waste generated by the conventional protocol (**Figure 1A**) and a “green” protocol (**Figure 3C**) incorporating all swaps proposed in this study. Both assume the user passages two cell lines per session. The conventional protocol is not intended to represent worst practice; we assume good sterile technique and reuse of serological pipettes where possible. In the sustainable protocol, T75 flasks are replaced with 10 cm dishes, waste reagents are decanted rather than aspirated with stripettes, and cells are passaged ratiometrically within the same vessel. Implementing these changes yields an estimated saving of 154 g plastic per culture session, and almost 15kg plastic annually. Large weights are shown in red, while smaller weights are shown in green.

| Action | Waste Stream | CO<sub>2</sub>e Use, g |  |  |  | Plastic Use, g |  |  |  |
| --- | --- | --- | --- | --- | --- | --- | --- | --- | --- |
|  |  | Single Swap | Weekly | Monthly | Annual | Single Swap | Weekly | Monthly | Annual |
| Conventional Protocol | No Disposal | 672 | 1345 | 5379 | 64542 | 187 | 374 | 1494 | 17933 |
|  | Incineration with E Recovery | 877 | 1754 | 7015 | 84184 |  |  |  |  |
|  | Autoclave + Incineration with E Recovery | 940 | 1881 | 7523 | 90281 |  |  |  |  |
|  | Clinical Waste | 1152 | 2305 | 9219 | 110627 |  |  |  |  |
| New Protocol | No Disposal | 125 | 250 | 998 | 11977 | 33 | 66 | 265 | 3177 |
|  | Incineration with E Recovery | 160 | 321 | 1283 | 15398 |  |  |  |  |
|  | Autoclave + Incineration with E Recovery | 172 | 343 | 1373 | 16478 |  |  |  |  |
|  | Clinical Waste | 210 | 420 | 1678 | 20142 |  |  |  |  |
| Savings by Updating Protocol | No Disposal | 548 | 1095 | 4380 | 52565 | 154 | 307 | 1230 | 14756 |
|  | Incineration with E Recovery | 717 | 1433 | 5732 | 68787 |  |  |  |  |
|  | Autoclave + Incineration with E Recovery | 769 | 1538 | 6150 | 73803 |  |  |  |  |
|  | Clinical Waste | 943 | 1885 | 7540 | 90485 |  |  |  |  |

### Swap 2: Use the Most Appropriate Culture Vessel

As shown in **Figure 1C**, selecting the most appropriate culture vessel is a straightforward yet effective way to reduce resource use and waste. Smaller flasks and dishes require fewer materials and generate less plastic waste, while still supporting most experimental needs. Depending on your experimental needs, consider whether transitioning to smaller flasks might be a viable option. Alternatively, T75 flasks and 10 cm dishes offer comparable surface areas, yet switching from a T75 flask to a 10 cm dish saves nearly 40 g of plastic and 174 g CO_2_e per use, equating to over 7 kg of plastic savings per researcher annually. Such adjustments can substantially reduce a laboratory’s plastic and carbon footprint without compromising experimental quality or flexibility.

### Swap 3: Passage Cells Within the Same Container

Another sustainable practice is to passage cells within the same culture dish or flask rather than transferring them to a new vessel. This strategy reduces the number of containers required, thereby lowering plastic consumption and conserving resources. As illustrated in **Figure 1D**, after passaging, a fraction of cells can be reseeded directly into the original vessel, rather than discarding it. In the laboratories where this approach was implemented, sterility was maintained simply by keeping the vessel closed between passages and only opening it within a clean biosafety cabinet; this ensured that container reuse did not compromise culture quality or introduce contamination.

By optimising vessel reuse in this way, laboratories can substantially decrease their reliance on single-use plastics while maintaining cell health and culture quality. For example, if a researcher replaces each culture vessel once per month (i.e., every eight passages), this would save approximately 14 culture dishes per month, or 168 dishes over a 48-week year. For T75 flasks, this equates to more than 9 kg of plastic and 44 kg CO_2_e savings annually per researcher.

### Swap 4: Replace Single-Use Haemocytometers with Reusable Alternatives

Switching to a reusable haemocytometer is another small change that yields substantial long-term impact. Each disposable haemocytometer generates 5 g of waste, without including packaging. This may seem negligible, but over time, this waste adds up significantly in busy labs. A researcher culturing two cell lines and passaging twice per week would generate nearly 350 g plastic (and almost 1.5 kg CO_2_e) annually from disposable haemocytometers alone. Adopting a reusable equivalent eliminates this waste and requires only routine cleaning between uses. In the laboratories we observed, the reusable haemocytometers were simply sprayed with 70% ethanol and wiped between counts, providing an effective and low-effort method for ensuring they remained contamination-free.

### Swap 5: Passage Cells Ratiometrically Instead of Centrifuging and Counting

Many laboratories include a centrifugation step to remove dissociation enzymes after cell detachment. However, once the enzyme is neutralised with fresh medium, this step is often unnecessary. Routine passaging can instead be achieved by simply diluting cells to the desired ratio in new medium. As shown in **Figure 1F**, this adjustment not only streamlines the workflow but also avoids the use of a 15 mL centrifuge tube, a p20 filter tip, and a disposable haemocytometer per passage, amounting to over 1.5 kg of plastic and 6.3 kg CO_2_e savings annually for a typical researcher.

### Swap 6: Consider which Waste Stream is Appropriate for Cell Culture Plastic

In many research laboratories, all cell culture waste is routinely classified as hazardous, requiring autoclaving followed by incineration with energy recovery. Increasingly, however, institutions are adopting on-site chemical decontamination protocols to inactivate biological material prior to disposal. When doing so, it is important to select disinfectants that are both effective and non-hazardous to the environment, thereby avoiding the transfer of sustainability burdens to other stages of the workflow.

This approach enables the diversion of a large proportion of cell culture plastics into the offensive waste stream, which is less energy-intensive and more cost-effective. It is recognised, however, that waste disposal requirements may differ when genetically modified organisms (GMOs) are present; in such cases, laboratories should consult local biosafety or health-and-safety officers to ensure compliance with institutional and regulatory frameworks.

When implemented appropriately, these practices maintain biosafety while reducing both greenhouse gas emissions and disposal costs. For example, under our standard protocol, switching from autoclaving plus incineration to chemical decontamination plus incineration with energy recovery would save approximately 6 kg CO_2_e per researcher annually. While incineration remains the standard end-point for decontaminated lab plastics, emerging recycling schemes are beginning to offer more circular alternatives. As these systems become more accessible, they may provide lower-carbon, lower-cost disposal routes for decontaminated lab waste.

### Other Considerations

In addition to consumable swaps, several broader practices can further reduce the environmental footprint of cell culture.

#### Biosafety Cabinet Use

Efficient operation of cell culture biosafety cabinets can substantially reduce both energy consumption and environmental impact. When left running continuously, these cabinets may consume more than half the energy of an average household (NIH Environmental Management System, 2021), highlighting the importance of switching them off whenever they are not in use. Regular maintenance is also essential to ensure that airflow grills remain unobstructed, as blockages not only increase energy demand but also elevate the risk of contamination by drawing in unfiltered air (Tay, 2020). The necessity of ultraviolet (UV) sterilisation should also be critically assessed: chemical disinfection is generally sufficient for routine operations (Mizuno et al., 2023), and where UV sterilisation is employed, shorter exposure periods are typically adequate (Meechan & Wilson, 2006). Energy use can be further reduced by turning off the cabinet’s fluorescent lighting, which accounts for up to 8% of total consumption (Phillips, 2009). Collectively, these practices can lower energy demand while maintaining sterility and safety in culture workflows.

#### Cold Storage

Ultra-low temperature freezers represent a major source of laboratory emissions. Raising freezer temperatures from −80 °C to −70 °C is supported by literature and community-shared data showing that sample integrity is maintained, and has the added benefit of reducing freezer strain, extending lifespan, and lowering running costs (Farley et al., 2015; University of Colorado Boulder Green Labs Program, 2011). Temperature fluctuations, rather than absolute temperature, have been shown to cause sample degradation (Mitchell et al., 2005), and freezers set at -80°C have greater fluctuations each time the door is opened (Farley et al., 2015). Proper storage practices, such as placing critical samples furthest from the door of the freezer, and packing available space with insulators, can offer more protection than simply setting a colder temperature. Additionally, where applicable, replacing mechanical -150 °C freezers with liquid nitrogen storage is a high-impact swap. Investigations at the Cancer Research UK Manchester Institute estimated that a -150 °C freezer emits more than 100 times the CO_2_e per sample compared with liquid nitrogen storage.

#### Animal-Derived Serums

The continued reliance on animal-derived serums in cell culture presents well-recognised ethical, environmental, and reproducibility challenges (Jochems et al., 2002; Van Der Valk et al., 2004; Wali et al., 2024). Foetal bovine serum is collected from bovine foetuses during slaughter, raising substantial animal welfare concerns. One potential alternative is donor bovine serum, where donor cattle are raised in a specific, controlled herd solely for blood donation, and blood is collected multiple times. As well as avoiding the ethical issues associated with foetal harvesting, donor bovine serum is available in larger volumes and is considerably less expensive than foetal bovine serum. Similarly, in contexts where horse serum (HS) is required, donor HS, collected non-lethally from living animals under veterinary supervision, offers a more ethically acceptable option. Across all serum types, reduction of serum concentration (Van Der Valk et al., 2010; Wali et al., 2024) or the adoption of serum-free (Jochems et al., 2002; Van Der Valk et al., 2004) and chemically defined (Utrecht University, 3Rs Centre Utrecht, 2025) where feasible represents a practical strategy to minimise reliance on animal slaughter while improving reproducibility.

#### Appropriate Reagent Volumes

Minimising the use of reagents such as culture media, trypsin, and PBS is a simple yet effective strategy to improve sustainability in cell culture. Sigma-Aldrich recommends using 0.5 mL of Trypsin per 10 cm^2^ of culture surface area, followed by neutralisation with twice the volume of complete growth medium (Sigma-Aldrich (Merck), n.d.). Adhering to these recommended volumes ensures effective cell detachment without unnecessary waste. Overfilling flasks with media or using excessive volumes of PBS for washing not only increases reagent consumption but also drives up associated plastic use. By calibrating reagent volumes vessel size and experimental purpose, researchers can significantly reduce both environmental impact and operational expenses, without compromising experimental quality.

#### Antibiotic use

Routine supplementation of culture media with antibiotics can mask suboptimal aseptic practice, compromise reproducibility, and contribute to avoidable chemical waste. Moreover, the large-scale use and disposal of antibiotic-containing media introduces pharmaceutically active compounds into laboratory waste streams, potentially contributing to antimicrobial resistance and environmental contamination (Kümmerer, 2009; Leung et al., 2012). Restricting antibiotic use to essential applications therefore reduces chemical burden, aligns with global efforts to minimise antimicrobial pollution, and promotes more rigorous aseptic technique, ultimately improving both sustainability and experimental reliability. In practice, this is typically achieved by simply omitting antibiotics from routine culture medium and maintaining strict sterile workflow—for example, working consistently within a biosafety cabinet, wearing a clean lab coat, and using clean gloves to minimise the risk of contamination.

### A More Sustainable Protocol

A more sustainable, “green” protocol incorporating all the steps described in this paper is illustrated in **Figure 3A**. As shown in **Table 5**, this updated workflow generates just 33.1 g of plastic across 5 items per culture session (culturing two lines simultaneously), compared with 186.8 g and 16 items under the conventional approach. Crucially, these adjustments do not compromise sterility or experimental quality: cultures are maintained to the same standard while avoiding unnecessary consumables. As shown in **Figure 3B** and **Figure 3C**, by replacing T75 flasks with 10 cm dishes, decanting reagents instead of aspirating with stereological pipettes, and passaging cells ratiometrically within the same vessel, researchers can save 154 g plastic per session—equivalent to almost 15 kg of plastic and 74 kg CO_2_e per year for a single user. Even greater reductions are possible when laboratories coordinate reference cell line maintenance (Swap 1). These data highlight that meaningful sustainability gains are possible without sacrificing scientific rigour. We encourage researchers to identify swaps relevant to their workflows and to use the accompanying tools and tables to estimate the associated reductions in plastic use and CO_2_e emissions.

**Table 5:** Estimated plastic generated per cell passaging session following the standard protocol (outlined in **Figure 3C**). Calculations assume one researcher passaging two cell lines in a single session. One cell culture session requires 5 disposable plastic items with a combined mass of 33.1 g, broken down by consumable type as shown. Large weights are shown in red, while smaller weights are shown in green.

| Consumable | Labels on Diagram | Consumable Weight, g | Number Required | Total Weight, g |
| --- | --- | --- | --- | --- |
| 10ml stereological pipette | C,D,E | 8.3 | 3 | 24.8 |
| 5ml stereological pipette | A | 7.5 | 1 | 7.5 |
| P1000 Filter Tip | B | 0.8 | 1 | 0.8 |
| Totals |  |  | 5 | 33.1 |

## CONCLUSION

This study demonstrates that routine cell culture workflows can be redesigned to substantially reduce plastic use and associated CO_2_e emissions without compromising experimental practice. Using an online calculator to provide a consistent framework for protocol-level environmental quantification, we show that relatively simple changes, including coordinating culture schedules, optimising vessel choice, passaging within the same container, and reusing appropriate tools, can generate substantial cumulative savings when applied to high-frequency laboratory activities. These interventions are practical, flexible, and readily transferable across research settings, allowing researchers to adopt those most compatible with their own experimental requirements and laboratory context.

Although not quantified explicitly here because of variation in procurement systems and pricing across institutions and countries, many of the changes described are also likely to reduce financial costs. Reducing consumable use, improving energy efficiency where feasible, and adopting lower-impact alternatives to commonly used reagents may all contribute to long-term financial savings alongside environmental benefit.

Beyond the specific interventions evaluated, this work provides an open-source tool for quantifying the environmental consequences of laboratory workflow design. The online calculator offers an accessible implementation of this framework, enabling researchers to assess and compare the impacts of protocol changes within their own laboratory settings. In this way, the study contributes both a set of evidence-based recommendations for greener cell culture practice and a practical tool for extending this mode of analysis to other workflows.

Taken together, these findings support a shift from general awareness of laboratory sustainability challenges towards routine, quantitative evaluation of everyday practice. Embedding environmental considerations into experimental design should be regarded not as an optional addition, but as part of robust, responsible, and forward-looking research practice. Wider adoption of standardised reporting approaches and environmental assessment tools will be important for accelerating the transition towards more sustainable life-science research.

## REFERENCES

Environmental Impact Calculator. (2026). SPARKHub. https://consumablescalculator.sparkhub.eu/

Farley, M., McTeir, B., Arnott, A., & Evans, A. (2015). Efficient ULT Freezer Storage: An Investigation of ULT Freezer Energy and Temperature Dynamics (Social Responsibility & Sustainability). University of Edinburgh.

Jochems, C. E. A., Van Der Valk, J. B. F., Stafleu, F. R., & Baumans, V. (2002). The Use of Fetal Bovine Serum: Ethical or Scientific Problem? Alternatives to Laboratory Animals, 30(2), 219–227. 10.1177/026119290203000208

Kümmerer, K. (2009). Antibiotics in the Aquatic Environment – A Review – Part I. Chemosphere, 75(4), 417–434. 10.1016/j.chemosphere.2008.11.086

Leung, H. W., Minh, T. B., Murphy, M. B., Lam, J. C. W., So, M. K., Martin, M., Lam, P. K. S., & Richardson, B. J. (2012). Distribution, Fate and Risk Assessment of Antibiotics in Sewage Treatment Plants in Hong Kong, South China. Environment International, 42, 1–9. 10.1016/j.envint.2011.03.004

Meechan, P. J., & Wilson, C. (2006). Use of Ultraviolet Lights in Biological Safety Cabinets: A Contrarian View. Applied Biosafety, 11(4), 222–227. 10.1177/153567600601100412

Mitchell, B. L., Yasui, Y., Li, C. I., Fitzpatrick, A. L., & Lampe, P. D. (2005). Impact of Freeze-Thaw Cycles and Storage Time on Plasma Samples Used in Mass Spectrometry Based Biomarker Discovery Projects. Cancer Informatics, 1(1), 98–104.

Mizuno, M., Matsuda, J., Watanabe, K., Shimizu, N., & Sekiya, I. (2023). Effect of Disinfectants and Manual Wiping for Processing the Cell Product Changeover in a Biosafety Cabinet. Regenerative Therapy, 22, 169–175. 10.1016/j.reth.2023.01.009

NIH Environmental Management System. (2021, October). How Much Energy Do Your Biosafety Cabinets Use? NIH Environmental Management System. https://nems.nih.gov/Documents/Newsletter/2021/10_October_2021/Biosafety_Cabinets.pdf

Phillips, D. S. (2009). Biological Safety Cabinets: Energy Efficiency Saves Resources. Thermo Fisher Scientific / BIOforum Europe (GIT Verlag GmbH & Co.). http://tools.thermofisher.com/content/sfs/brochures/EnergyEfficiencySavesResources_BioforumE_0509.pdf

Ragazzi, I., Farley, M., Jeffery, K., & Butnar, I. (2023). Using Life Cycle Assessments to Guide Reduction in the Carbon Footprint of Single-Use Lab Consumables. PLOS Sustainability and Transformation, 2(9), e0000080. 10.1371/journal.pstr.0000080

Schneegans, S., Straza, T., & Lewis, J. (2021). UNESCO Science Report: the Race Against Time for Smarter Development. UNESCO Publishing: Paris.

Sigma-Aldrich (Merck). (n.d.). Trypsin Cell Dissociation Protocol [Protocol]. Retrieved 1 September 2025, from https://www.sigmaaldrich.com/GB/en/technical-documents/protocol/cell-culture-and-cell-culture-analysis/mammalian-cell-culture/cell-dissociation-with-trypsin

Tay, A. (2020, November 5). How Efficient Is Your Biosafety Cabinet? Lab Manager. https://www.labmanager.com/how-efficient-is-your-biosafety-cabinet-24221

University of Colorado Boulder Green Labs Program. (2011). Biological Samples Stored Long Term at - 70 C or Warmer [Data set]. https://docs.google.com/spreadsheets/d/136A8VQmOrWUFVP_EW3Q8wF4dNmRe5I9bmM6KkC8aH1o/edit?gid=702800635#gid=702800635

Urbina, M. A., Watts, A. J. R., & Reardon, E. E. (2015). Labs Should Cut Plastic Waste Too. Nature, 528(7583), 479–479. 10.1038/528479c

Utrecht University, 3Rs Centre Utrecht. (2025). Fetal Calf Serum-Free Database (SCR_018769) [Data set]. https://fcs-free.sites.uu.nl/database/

Van Der Valk, J., Brunner, D., De Smet, K., Fex Svenningsen, Å., Honegger, P., Knudsen, L. E., Lindl, T., Noraberg, J., Price, A., Scarino, M. L., & Gstraunthaler, G. (2010). Optimization of Chemically Defined Cell Culture Media – Replacing Fetal Bovine Serum in Mammalian in Vitro Methods. Toxicology in Vitro, 24(4), 1053–1063. 10.1016/j.tiv.2010.03.016

Van Der Valk, J., Mellor, D., Brands, R., Fischer, R., Gruber, F., Gstraunthaler, G., Hellebrekers, L., Hyllner, J., Jonker, F. H., Prieto, P., Thalen, M., & Baumans, V. (2004). The Humane Collection of Fetal Bovine Serum and Possibilities for Serum-Free Cell and Tissue Culture. Toxicology in Vitro, 18(1), 1–12. 10.1016/j.tiv.2003.08.009

Wali, M. E., Karinen, H., Rønning, S. B., Skrivergaard, S., Dorca-Preda, T., Rasmussen, M. K., Young, J. F., Therkildsen, M., Mogensen, L., Ryynänen, T., & Tuomisto, H. L. (2024). Life Cycle Assessment of Culture Media with Alternative Compositions for Cultured Meat Production. The International Journal of Life Cycle Assessment, 29(11), 2077–2093. 10.1007/s11367-024-02350-6

Weber, P. M., Michelsen, C., & Kerou, M. (2025). What’s in our Bin? Labs Kick off and Demand the Transition Towards a Circular Economy for Lab Plastics. EMBO Reports, 26(2), 297–302. 10.1038/s44319-024-00360-x

